# Bundle-specific fornix reconstruction for dual-tracer PET-tractometry

**DOI:** 10.1101/423459

**Authors:** Francois Rheault, Maggie Roy, Stephen Cunnane, Maxime Descoteaux

## Abstract

Tractography is known to have problems reconstructing white matter bundles that are narrow, have high curvature, or go through partial volume voxels contaminated by CSF or gray matter. One such bundle is the fornix, the major output tract of the hippocampus, which is especially problematic with aging. Hippocampal atrophy and ventricular expansion make the fornix even harder (often impossible) to track with current state-of-the-art techniques. In this work, a bundle-specific tractography algorithm is proposed to fully reconstruct the fornix. By injecting shape, position, and orientation priors, fornix reconstruction is markedly is improved. We report an increase in spatial coverage and better reproducibility across test-retest. These improvements over classical tractography algorithms also enable tractometry of the fornix to be combined with dual-tracer positron emission tomography (PET) data in participants with mild cognitive impairment (MCI). MCI participants underwent a multi-modal brain imaging before and after a 6-month daily ketogenic supplement. We report, for the first time, significant diffusion measures and 18F-fluorodeoxyglucose (FDG) uptake differences in specific sub-sections of the fornix after the ketogenic supplement.

## 1 Introduction

The fornix is a white matter (WM) bundle playing a vital role in memory and is therefore very relevant to analyze in cognitive decline associated with aging. The fornix originates from the hippocampi and join beneath the splenium of the corpus callosum to form the body of the fornix. The anterior body of the fornix then divides to enter both mamillary bodies. Reconstruction of the fornix white matter bundle for the tractography community is similar to hippocampus segmentation in the field of MRI segmentation (Kantarci, 2014), as it is an important tract of the memory structural network and thus, closely linked to neurodegenerative disease in aging population (Lister and Barnes, 2009).

In aged populations, hippocampal atrophy and ventricular expansion make the fornix extremely hard-to-track (Fletcher et al., 2013). Specific acquisition schemes, such as FLAIR-DTI (Chou et al., 2005) which removes the signal from the ventricles, have been developed to facilitate tractography of the fornix. However, existing databases, such as Alzheimer’s Disease Neuroimaging Initiative (ADNI) (Petersen et al., 2010), often use classical diffusion MRI (dMRI) acquisitions tailored for diffusion tensor imaging (DTI) processing (Sudlow et al., 2015; Marek et al., 2011). Alternatively, free-water (FW) elimination processing (Pasternak et al., 2009; Concha et al., 2005) can reduce CSF contamination causing partial volume effect around the ventricles and improve DTI tractography. Leading to an improved fornix reconstruction by typically increasing the spatial extent. However, FW-corrected DTI remains a single fiber modeling technique, which is known to be sub-optimal for tractography of bundles with high curvature and crossing fiber configurations (Girard et al., 2014; Tournier et al., 2008).

Without anatomical priors, researchers targeting difficult bundles of interest often use tractography with a brute force approach, which consists of an aggressive seeding strategy across the whole brain, often generating tens of millions of streamlines (Renauld et al., 2016), in the hope of having a few lucky streamlines surviving manual dissection (Catani et al., 2013) or automatic segmentation (Garyfallidis et al., 2017). The low quality and difficulty of fornix tractography reconstruction explains why this bundle is rarely reported in the literature, particularly in the aging population.

In the present study, a bundle-specific approach was developed to reconstruct the fornix, increasing the spatial extent and improving the reproducibility across time points and participants. This allowed a dual-tracer PET fornix tractometry application study on 23 participants with mild cognitive impairment (MCI) at two time points, before and after a 6-month ketogenic supplement of MCT (medium chain triglyceride) or placebo.

## 2 Methods

### 2.1 Multi-modal diffusion & PET imaging

MCI participants were randomized in the placebo (N=11) or MCT (N=12) groups. Participants underwent multi-modal brain imaging before and after a 6-month daily supplement of 30 g/day MCT (Croteau et al., 2017). The protocol consisted of an isotropic 1 mm T1-weighted (T1w) image, followed by a 1.8 mm isotropic high angular resolution diffusion imaging (HARDI; 60 directions, b=1500 s/mm^2^), and a blip-up/blip-down b=0 s/mm^2^ acquisition to correct for distortions. Diffusion-weighted images were upsampled to 1 mm isotropic resolution, diffusion tensor measures, free-water index (Pasternak et al., 2009), and fiber orientation distribution function (fODF) (Tournier et al., 2012; Descoteaux et al., 2009) were computed using Dipy (Garyfallidis et al., 2014).

The MRI protocol was followed by a dual PET tracer session: AcAc (11Cacetoacetate) first, followed by FDG (18F-fluorodeoxyglucose)(Croteau et al., 2017). SUV (standardized uptake values) summed-images from dynamic acquisitions were used. Finally, PET images were co-registered to the T1w image and then to upsampled dMRI data using ANTs (Avants et al., 2010). Average PET tracer uptake and tract-profiling along 10 sections of the fornix were measured. Data are expressed as delta (*Δ*) (post-supplementation minus pre-supplementation).

### 2.2 Fornix reconstruction

The bundle-specific tractography (BST) approach is similar to the one proposed in (Rheault et al., 2017) and is composed of three steps.

1. Build a template of streamlines that represents the shape and position of the fornix, the template must represented the full spatial extent of the fornix.
2. Build spatial anatomicals priors that represents the endpoints and the general position of the bundle (seeding and tracking mask). To solve partial volume effect difficulties below the ventricules, tracking was allowed in GM to maximize the chance of fully reconstructing the fornix.
3. Build the orientation priors from the track-orientation distribution of the template of streamlines, and then create the enhanced fiber orientation distrubtions (E-fODF).

### 2.3 Template creation

A population-specific template was created using *ANTs Multivariate Template Construction* (Avants et al., 2010) due to the major structural changes appearing with age. T1w images and the fractional anisotropy (FA) maps from the pre-supplementation acquisitions (n=23) were used as the input. The resulting average T1w and FA images can be seen in Figure 1A. To create an initial streamline-template of the fornix, classical probabilistic tractography (Tournier et al., 2012), within WM and GM, was used on the pre-supplementation acquisitions. Each whole brain tractogram was then automatically segmented by moving streamlines to a common template space and then ROIs segmentation was used. Since each participant had a poorly saturated fornix, meaning most had less than 50 streamlines and that were not covering much of the expected volume, streamlines were concatenated in the common template space, forming a single, more dense, fornix template, as seen in Figure 1 B.

**Fig. 1.**
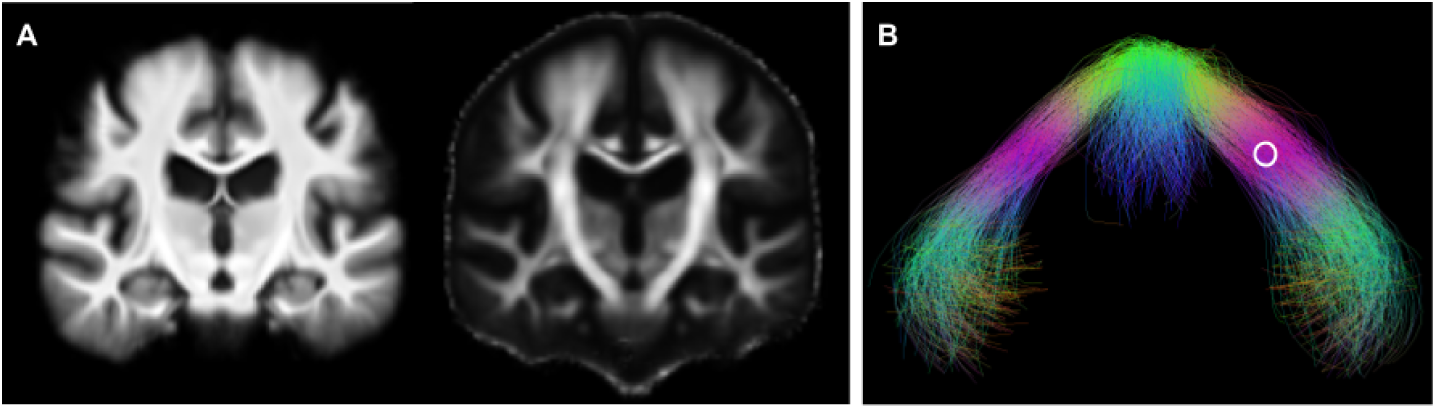
A) T1 and FA template from the 23 pre-supplementation acquisitions B) Representation of the average fornix for the 23 participants (zoomed). The white circle shows the region of origin of the fODF of Figure 2.

## 2.4 Orientation enhancement & mask extraction

The newly created fornix template was then used to inform and enhance fiber orientation distribution function (fODF) needed by tractography, ultimately to help reconstruct the full fornix of each participant. First, the streamline-template was deformed to a participant native diffusion space. Then, at each voxel, an orientation prior was created from the streamline-template to reinforce the appropriate directions (Rheault et al., 2017; Dhollander et al., 2014), as seen in Figure 2. Seeding and tracking masks were also adapted to improve computational performance by restricting the regions where tractography is allowed (Rheault et al., 2017; Renauld et al., 2016). Tissue types were modified in a region under the ventricles particularly prone to partial volume effect, typically identified as CSF or GM, to allow tractography to propagate through the body of the fornix to the back of the thalamus.

**Fig. 2.**
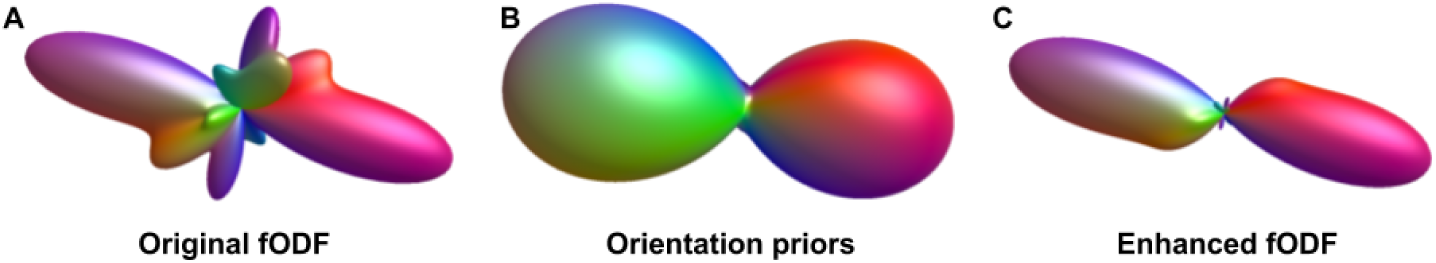
Application of the orientation prior in one voxel. The original fODF contains more than one fiber populations (A) and the orientation prior (B) reduces the influence of lobes that are not desirable or useful for the fornix reconstruction (C). The fODF shown is from the region within the white circle in Figure 1B.

## 2.5 Tractography

Anatomically-constrained particle filtering tractography (PFT) (Girard et al., 2014) with the bundle-specific probability maps and the bundle-specific enhanced-fODF was used to reconstruct the fornix (20 seeds per voxel, length threshold 40mm-160mm, and otherwise default parameters (Girard et al., 2014)). A classical approach was used as a comparison, i.e anatomically-constrained PFT with default parameters and no bundle-specific input (original fODF and probability WM-GM-CSF masks).

## 2.6 Bundle segmentation

To compare the fornix across all datasets, an automatic segmentation was designed using anatomical landmarks in the population-specific template to reduce bundle segmentation variability. The thalamus and hippocampus were obtained with *Freesurfer* sub-cortical segmentation (Desikan et al., 2006) and mammillary bodies were manually delineated. To be considered valid, streamlines had to end in either the hippocampi or the mammillary bodies and had to pass through the body of the fornix (Catani et al., 2013). The thalamus were used as an exclusion region of interest because the fornix has to curve around it. The anatomical regions of interest used for the segmentation can be seen in Figure 3.

**Fig. 3.**
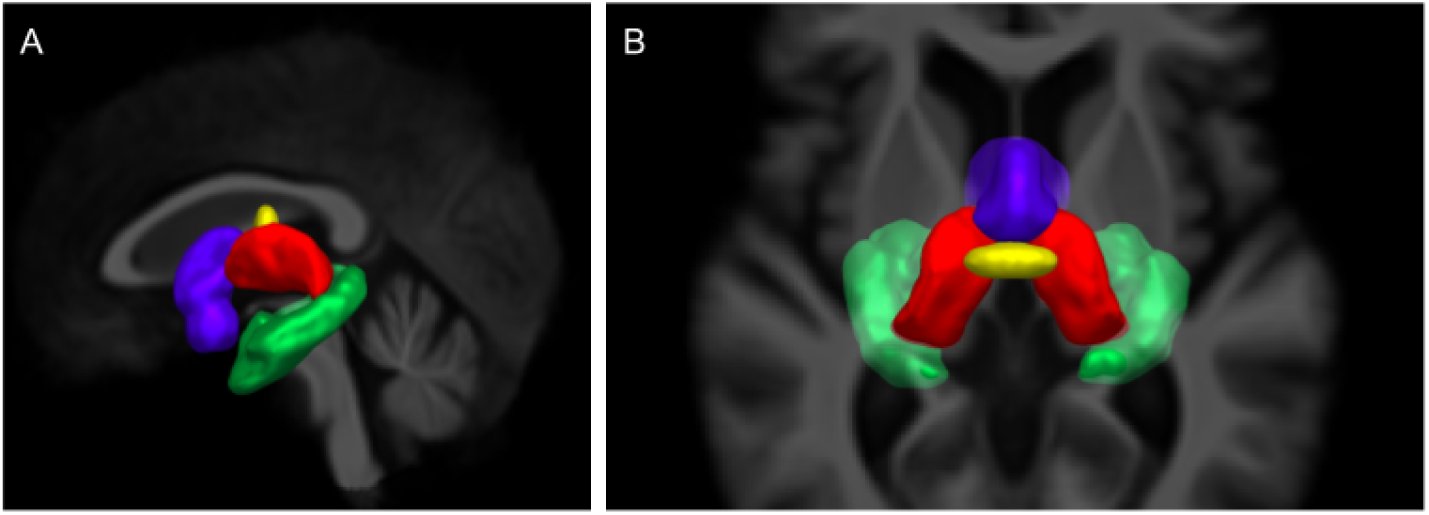
Sagittal view (A) and axial view (B) of the regions of interest used for the fornix segmentation in template space. Mammillary bodies in blue, body of the fornix in yellow, both thalami in red and both hippocampi in green.

## 3 Results

### 3.1 Tractography

To evaluate fornix reconstructions, the spatial extent of the whole fornix was computed for each methods (classical vs enhanced), for each time point (pre vs post-supplementation) and each MCI group (placebo vs MCT). As seen in Figure 4A, the classical method performs poorly in terms of volume against the enhanced version. Overall, the fornix reconstructed with the addition of priors were more dense and had a more coherent shape, as seen in Figure 5.

**Fig. 4.**
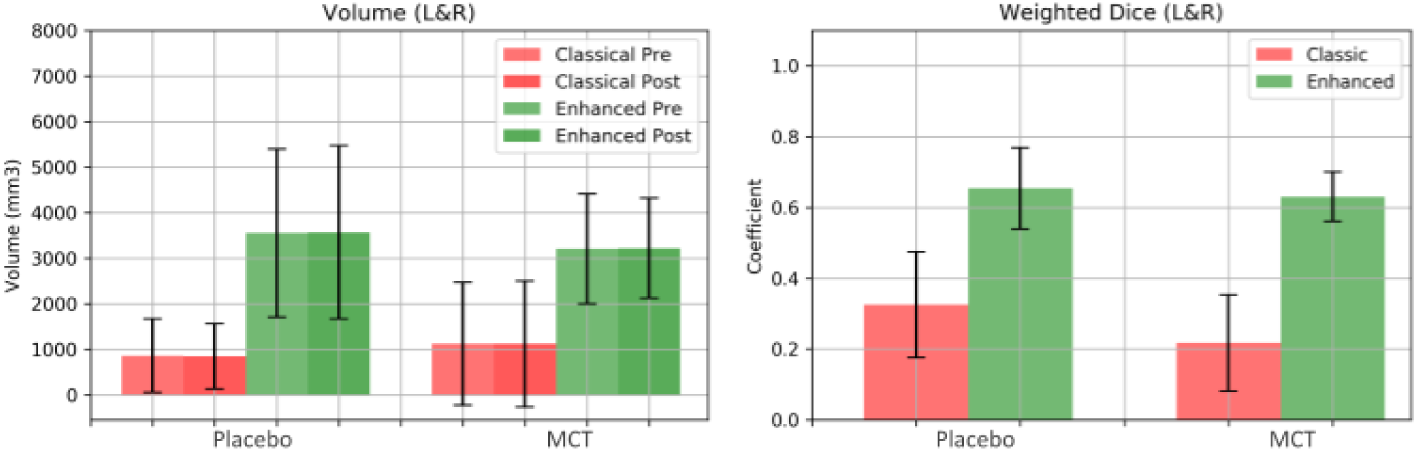
A) Average volume of the whole fornix comparison across acquisitions and groups for both methods (classical in red and the enhanced method in green). B) Weighted dice coefficient showing average overlap across time points for both groups. Data are expressed as mean and standard deviation.

**Fig. 5.**
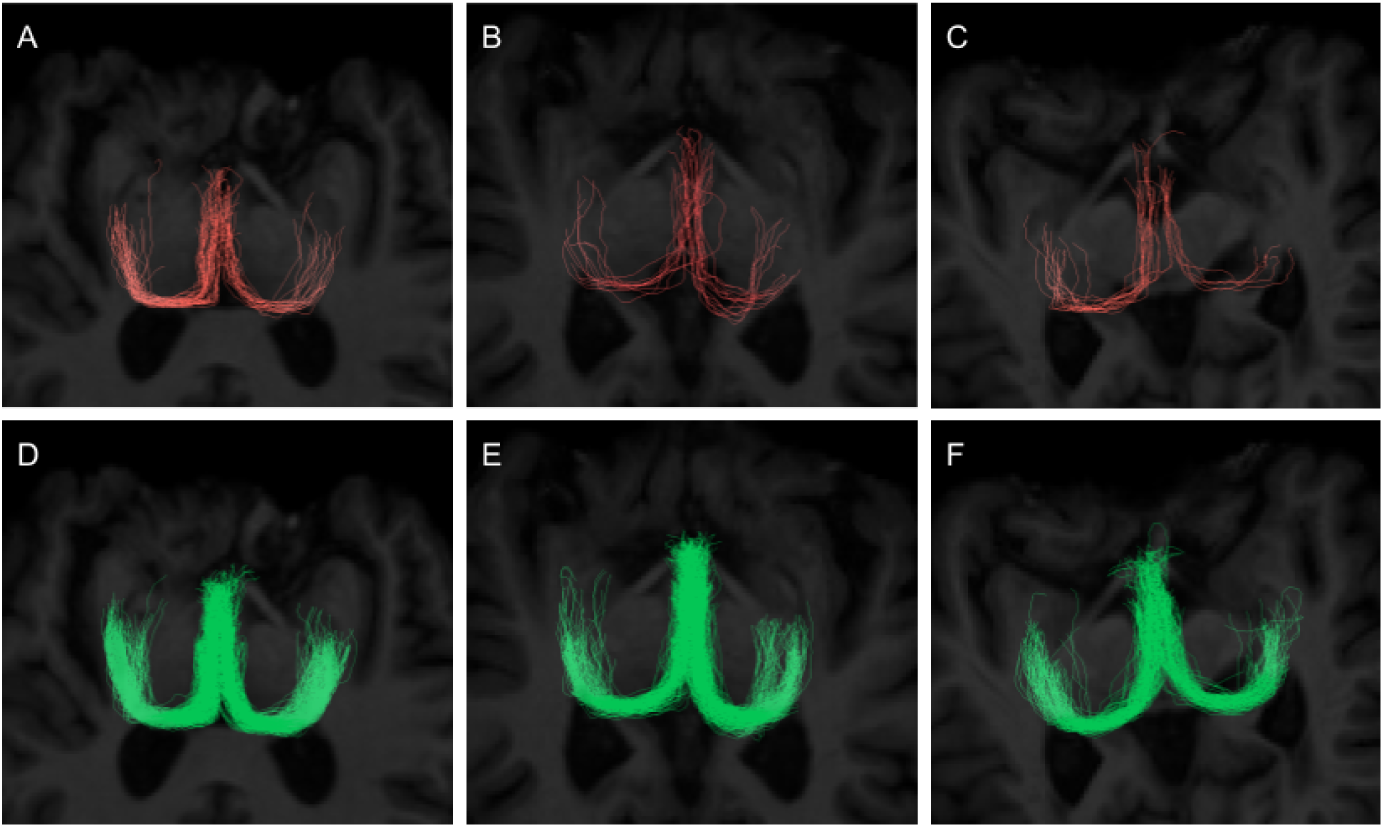
Comparison of the tractography output for 3 participants (presupplementation) between the classical method (A, B, C) and the enhanced method (D, E, F).

Using the two time points as a test-retest datasets for each participant, overlap was measured using weighted Dice (Cousineau et al., 2017) to verify the degree of reproducibility of each fornix reconstruction. As seen in Figure 4B, the classical method has severe difficulties to reconstruct the bundle across acquisitions, while the enhanced version obtains better overlap.

Overall, both the volume and weighted-Dice measures need to be taken into account together because the classical method obtains a low volume and low overlap, while the enhanced method obtains a higher volume and higher overlap, both which are needed for robust tractometry (Cousineau et al., 2017).

### 3.2 Tractometry

A preliminary tractometry analysis (Cousineau et al., 2017) was then performed on the enhanced fornix reconstructions using both the diffusion and dual-tracer PET data. Average PET tracer uptake and tract-profiling along 10 sections (5 in the left part, 5 in the right part) of the fornix were measured, as seen in Figure 6. Data are expressed as delta (*Δ*) (post minus pre-supplementation). All DTI measures (FA, mean, axial, and radial diffusivities), as well as free-water index were also submitted to the same tractometry analysis. First, lower *Δ*FDG uptake was observed in the tract-profiles of the MCT group. Sections showing statistically significant differences are located in the left part of the body (sections 3, 4 and 5; *p*= 0.05, 0.01 and 0.03 respectively) as illustrated in Figure 6. Tract-profiling of *Δ*AcAc uptake showed no statistical difference with the current number of participants.

**Fig. 6.**
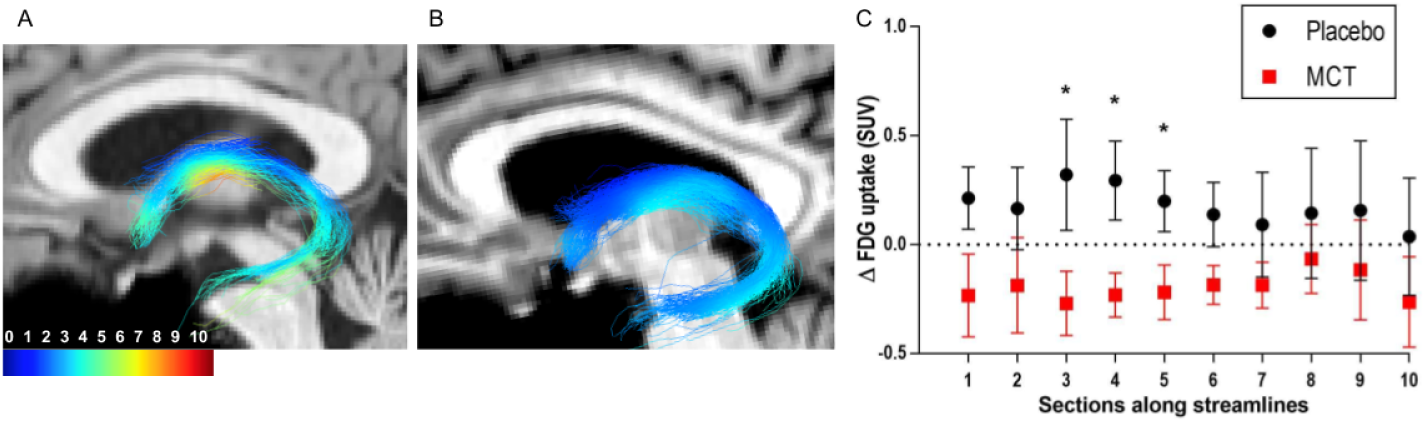
Sagittal view of the fornix from one MCI participant in the placebo group (A) and MCT group (B) at post-supplementation. The color scale represents 18Ffluorodeoxyglucose (FDG) uptake in SUV (standardized uptake values) along streamlines. The fornix was subsampled in 10 sections: sections 1-5 are located on the left part starting at the hippocampus; sections 6-10 are located in the right part ending at the hippocampus. C) FDG uptake along the 10 subsections of the fornix. FDG uptake was significantly lower for the MCT group in sections 3, 4 and 5. Data are expressed as delta (post- minus pre-supplementation) and mean *±SEM.*

In Figure 7, we show tract-profiles for *Δ*free-water index, mean and radial diffusivity; higher values in the MCT group were found in the left part of the fornix (sections 2 and 3) for the free-water and mean/radial diffusivities. When PET and diffusion data were averaged for the whole fornix, no statistically significant differences were observed post-intervention between the placebo and MCT groups.

**Fig. 7.**
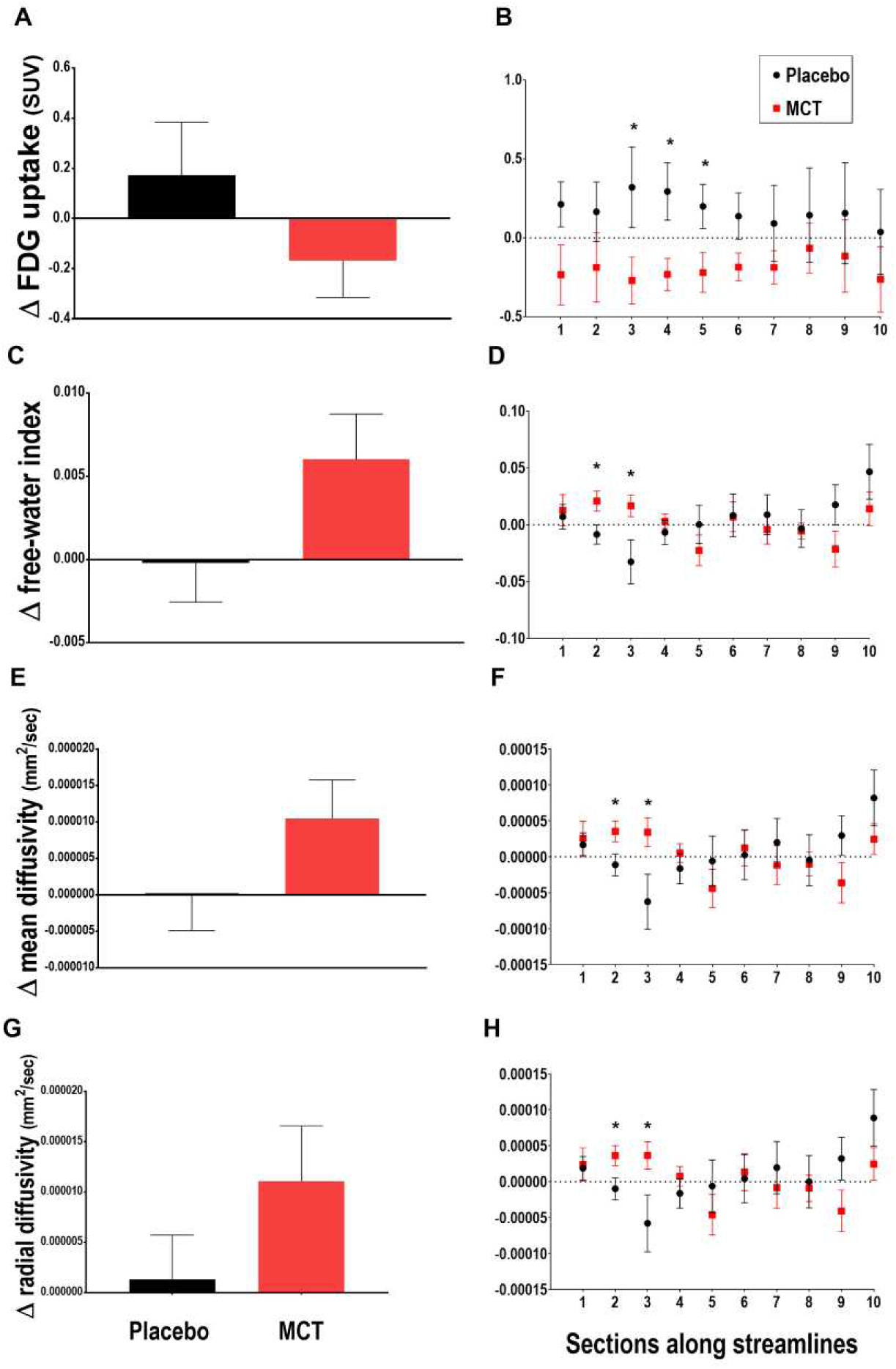
Average (left) and tract-profiles (right) for FDG PET and diffusion measures (free-water index, mean and radial diffusivities) in MCI participants of the placebo and MCT groups. The fornix was subsampled in 10 sections: sections 1-5 are located on the left part starting at the hippocampus; sections 6-10 are located in the right part ending at the hippocampus. When PET and diffusion data were averaged for the whole fornix (A, C, E, G), no statistically significant difference were observed between the placebo and MCT groups, whereas sections 2 and 3, on the left side of the fornix, show statistical significance for free-water index (D), mean (F) and radial diffusivities (H). Data are expressed as delta (postminus pre-supplementation) and mean SEM.

## 4 Discussion

### 4.1 Quality of the reconstruction

In this aged cohort, it is important to note that the fornix reconstruction with classical HARDI-based tractography methods was either impossible (complete fornix being un-trackable) or the the quality so poor that a tractometry study was impossible. Hence, a dedicated bundle-specific fornix strategy was necessary.

In this work, quality of a tractography reconstruction is evaluated solely on spatial extent and reproducibility after a strict segmentation. This may not reflect the true anatomy of the fornix, as the full spatial extent remains unknown and was limited by our image resolution. On average, the test-retest Dice score was 0.32 ± 0.15 for classical tractography and 0.65± 0.11 for the proposed approach. Further improvements are needed to reach Dice scores of 0.8 or above. Seeding along the fornix instead of seeding from the endpoints could potentially produce an increased in the number of valid streamlines. Moreover, the initial choice of tracking direction could also be improved by selecting the largest fODF lobe, which is the one that represents the main fiber population of the fornix due to the orientation priors.

One of the major challenges for fornix-tractography is to obtain an adequate WM/GM/CSF segmentation. For this work, *FSL FAST* was used on the T1w images. On numerous occasions, partial volume effect due to enlarged ventricles was observed and WM was mis-classified as GM. More robust approaches using advanced tissues segmentation algorithms and priors suited for the aging brain are needed for anatomically constrained tractography.

It is also important to note that the creation of a population-specific template (T1, FA) was useful to facilitate the registration to a common space for analysis. Using a template such as MNI152, (Fonov et al., 2011) composed mainly of healthy participants, would have produce mediocre registration due to the atrophy of the cortex and enlarge ventricles in an older cohort (Avants et al., 2010).

### 4.2 Alternative approaches to reconstruct the fornix

Other dMRI acquisitions or modeling could help tractography to accurately reconstruct the fornix. As mentioned earlier, FW elimination through modeling or through FLAIR-DTI can help reduce partial volume effect caused by the CSF around the ventricles and help to improve tractography (Chou et al., 2005). However, the required acquisition is time consuming and most available databases do not use such dMRI sequences. Furthermore, the local model remains single-fiber based and comes with its own downside when other bundles of interest are targeted.

Another potential technique that could improve tractography of the fornix is local modeling approaches computed with multi-tissue response functions (Jeurissen et al., 2014). This could reduce the impact of CSF contamination around the ventricles. This local modeling technique requires acquisitions with multiple b-values, but the work by (Dhollander and Connelly, 2016) recently proposed a multi-tissue local model that only requires single shell acquisition, which seems promising.

### 4.3 Dual-tracer PET tractometry

Tract-profiling showed differences in FDG uptake between the two groups, which were not found when only the average FDG uptake in the fornix was compared. FDG uptake near the body and more left sections, seemed more affected by the MCT supplementation compared to right sections of the fornix. Therefore, the dense part of bundles may be more susceptible to changes in fuel (glucose) uptake. To our knowledge, this is the first report of FDG uptake in any WM fascicles in humans. The assessment of WM FDG uptake using PET enables the evaluation of WM energy metabolism *in vivo*. WM energy supply (axonal and oligodendrocytes), which serves mainly for resting potentials, myelin synthesis and intracellular trafficking of molecules, is crucial to sustain adequate axonal function (Harris and Attwell, 2012; Wender et al., 2000) and may be linked to the pathogenesis of MCI. Finally, it is critical to re-emphasize that this tractometry analysis would not have been possible without our enhanced bundle-specific fornix tracking. In most participants, the fornix was un-trackable or just composed of 10 streamlines or less in the mid-body of the fornix.

### 4.4 Future work

Tractometry, using dual-tracer PET measures, along the fornix is now possible. Insightful measurements can be extracted using tractometry, which would have been missed by averaging of measures done on the full bundle. Future work will include a tractometry analysis using dual-tracer PET to assess whether a ketogenic supplement has impact on fuel uptake in other bundles, such as the posterior cingulum of MCI participants. Cerebral metabolic rate of FDG and AcAc will also be computed to compare quantitative measures between groups and this work will be extended to more participants and combined with grey matter analysis. A second objective is to map changes along the length of the fornix using various PET metric in healthy controls, MCI and Alzheimer’s patients to further evaluate the role of WM connections in cognitive impairment.

## 5 Conclusion

Using a specific fornix streamline-template, position and orientation priors were injected to tractography in order to overcome fornix reconstruction difficulties in the aging brain. This process greatly improves the spatial coverage and reproducibility of tractography of the fornix in MCI. Without this tractography improvement, tract-based fornix investigation was simply impossible. Bundle-specific tractography could have a positive impact on aging studies using diffusion MRI to analyze specific WM fascicles.

